# Chameau (HBO1) regulates starvation in a temperature-dependent manner in *Drosophila melanogaster*

**DOI:** 10.1101/2025.01.17.633563

**Authors:** Anuroop Venkateswaran Venkatasubramani, Toshiharu Ichinose, Ignasi Forne, Hiromu Tanimoto, Shahaf Peleg, Axel Imhof

**Author notes:** Max-Delbrück-Centrum für Molekulare Medizin (MDC), Robert-Rössle-Str. 10, Building 31.2, 13125 Berlin, Deutschland.

## Abstract

The body temperature of *Drosophila melanogaster* depends on the extrinsic temperature. Numerous studies in fruit flies have shown that environmental temperature has an effect on metabolism, lifespan and starvation resilience. We have previously shown that Chameau (Chm), a MYST-domain acetyltransferase, promotes aging but also increases starvation resilience. As starvation resilience is highly temperature dependent, we explored the effect of temperature on starvation resilience in fruit flies with reduced *chm* expression. Strikingly, we found that an increase of 2°C was sufficient to restore starvation resilience in *chm* mutants. The increase in temperature rescued the dampened expression of genes involved in insulin, hormone and starvation response as well as a reduced rate of weight loss and misregulation of trehalose, which we observed in *chm* mutants at 23°C. Our data show that whereas Chm has an important role in regulating starvation at 23°C and below, it becomes obsolete at higher temperatures. Our finding that a gene plays an important role only under specific environmental conditions has important implications in light of the recent global change of climate conditions.

## Introduction

One of the paramount environmental factors that shapes the behavior of the organisms is temperature. Change in temperature affects many physiological processes such as basal metabolic rate, development, longevity and survival, especially in cold-blooded organisms (**Pijpe et al., 2007**). Insects, fish and nematodes are poikilothermic organisms whose body temperature changes with the environmental conditions and therefore have developed various adaptations to react to alterations in the environment **(Fast et al., 2017; Mołoń et al., 2020**). In a constantly changing environment, adaptation to such novel conditions becomes key for survival. Therefore, poikilothermic organisms can change their behavior or physiology in response to even very subtle environment changes (**Jang & Lee, 2018; Karan & David, 2000; Padmanabha et al., 2011; Pijpe et al., 200**7).

One such organism that has populated most habitats of the world is the fruit-fly, *Drosophila melanogaster*. Multiple studies have shown the influence of temperature on its development, lifespan, metabolism and physiology. For example, increase in temperature accelerates development, metabolism and aging in fruit flies (**Goh et al., 2021; Lin et al., 2023; Mołoń et al., 2020**). Despite the fact that it has been shown that even subtle changes, such as an increase of 2-3°C can have a substantial effect on fly physiology (**Mołoń et al., 2020**), most studies that address the molecular pathways involved in adaptation focus on relatively large changes in conditions, such as heat or cold shock.

We have recently shown that reduction of the acetyltransferase Chameau (Chm) results in an impaired starvation response when measured at 23°C. We have also detected that mutants of *chm* display substantial changes in expression of metabolic genes, proteins and the post-translational modification of proteins affecting physiology and starvation resilience (**Venkatasubramani et al., 2023**). Interestingly, population genetic studies in *Drosophila* identified *Chm* as one of the chromatin factors that are differentiated between tropical and temperate populations of *Drosophila melanogaster*. The *chm* gene region contains single nucleotide polymorphisms (SNPs) in varying densities depending on the ambient conditions of the fly population (**Croze et al., 2017; Levine & Begun, 2008**). Accordingly, we were wondering whether the role of Chm in mediating starvation response is sensitive to differences in environmental temperatures. Notably, our lab has already shown that exposure of fruit flies to different temperatures results in altered activity of enzymes involved in citrate synthesis and acetyl-CoA and correspondingly, changes in those metabolites and downstream histone acetylation (**Peleg et al., 2016**). These findings raise the possibility that Chm may play a selective role in the adaptation and survival upon temperature change.

To explore this, we decided to assess the effects of temperature on *chm* mutants and its ability to survive starvation. Our data show that the starvation phenotype of *chm* mutants was indeed temperature specific. We also observed an effect of temperature on the molecular changes observed in *chm* mutant fly strains such as the changes in histone PTMs, the transcriptome and the metabolome. Our observations confirm the role of Chm as an enzyme that mediates adaptation to stress under very specific environmental conditions.

## Results

### Mild temperature changes affect starvation response in chameau mutant flies

To test the influence of temperature on starvation resilience, we raised flies within a temperature range from 18-27*°*C. As previously shown, *chm^MYST/+^* flies show impaired survival upon starvation at 23*°*C (**Venkatasubramani et al., 2023**), which was even larger at lower temperatures. However, at temperatures of 25*°*C and above, the phenotype was no longer observed (**Fig. 1A-B and Supplementary Fig. 1A-D**). Although male and female fruit flies have differences in starvation susceptibility (**Bhakti Chandegra et al., 2017; Lin et al., 2023**), they show a similar temperature dependency (**Fig. 1C-D and Supplementary** Fig. 2A-D), which we even observed when flies were only partly kept at higher temperature (**Supplementary Fig. 3**). This was independent of geographic location as similar results were observed from two different labs in Germany and Japan (**Supplementary Fig. 4A-B; see Fly maintenance and Starvation assay sections in Materials and Methods**). We also observed a similar phenotype in a *chm* RNAi line (*chm^RNAi^*, crossed with *da-Gal4)* in both males and females, further validating temperature dependent differences in starvation resilience of flies with a reduced *chm* expression (**Fig. 1E, Supplementary Fig. 5A-B**). Taken together, we concluded that the impaired starvation in *chm* mutants compared to control flies is temperature dependent and is manifested only at colder temperatures.

**Figure 1:**
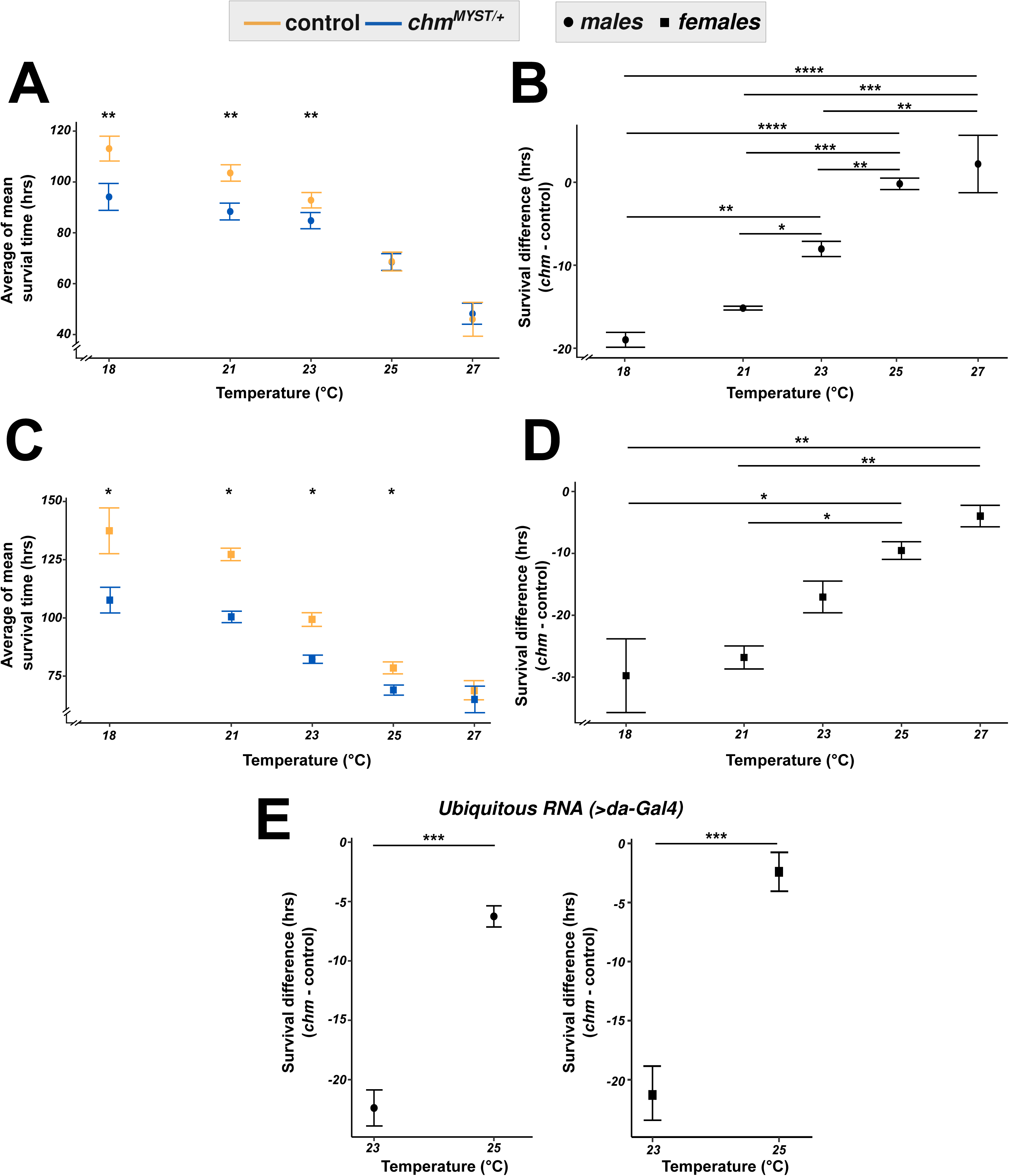
Temperature changes affect the starvation response in chm mutant flies irrespective of the gender. A) Average mean survival time in males and C) Average mean survival time in females of control and *chm^MYST/+^* mutants upon starvation at temperatures of 18°C (N = 3, paired), 21°C (N = 3, paired), 23°C (paired data obtained from **Venkatasubramani et al., 2023**), 25°C (N = 5 [males] and 4 [females], paired) and 27°C (N = 3, paired) B) Survival difference in males and D) Survival difference in males (difference in mean survival time in hours between mutant and control) from A) and C) respectively E) Survival difference (difference in mean survival time in hours between mutant and control) in male (left) and female (right) flies at 23°C (paired data obtained from **Venkatasubramani et al., 2023**) and 25°C (N = 4, paired) in *chm^RNAi^* flies as compared to its corresponding control *Non-significant values are not shown (* - p < 0.05, ** - p < 0.01, *** - p < 0.001, **** - p < 0.0001). Each replicate had at least 100 flies and paired t-test with FDR correction was performed for data in A), C) and E, while ANOVA followed by Tukey test correction was performed for data in B) and D). Error bars represent SEM*

As fruit flies have been shown to consume food differently when grown at different temperatures (**Klepsatel et al., 2019**), we measured the food consumption in control and *chm^MYST/+^* male flies at temperatures of 23*°*C and 25*°*C. Food intake and excretion showed no differences at 23*°*C and a marginal reduction in *chm^MYST/+^* flies at 25*°*C, which was not statistically significant (**Supplementary Fig. 6A-B**). Overall, the total food consumption showed negligible differences irrespective of both genotype and temperature (**Supplementary Fig. 65C**).

### Profiling of histone post-translation modifications indicates the role of acetylation in genotype, temperature and starvation response

Chm is an acetyltransferase that has been shown to affect H3 and H4 acetylation (**Feller et al., 2015; Nakagawa et al., 2015**). Therefore, we compared the histone modification levels of control and *chm^MYST/+^*male flies maintained at 23°C and 25°C under fed and starved conditions (**Dataset 1**). We combined the data into early (0-10 hours) and late (24-48 hours) starvation time points. A principle component analysis of H4 acetylation modification patterns revealed a distinct separation based on both temperature and genotype (**Fig. 2A**). Among the modifications that contributed the most to the separation were H4K12ac and H4 di-acetylation, which are bonafide targets of Chm (**Fig. 2B**) (**Feller et al., 2015**). While H4K12ac was significantly lower in *chm^MYST/+^* flies, H4 di-acetylation was also affected by temperature. Furthermore, the starvation induced changes of these modifications differed depending on the temperature (**Fig. 2C-D**). In contrast to the H4 acetylations, the acetylations in H3 showed only minor changes in *chm^MYST/+^* flies and mainly responded to a change in temperature (**Fig. 2E-H**). H3 methylation did not show any clustering, indicating the minor contribution of H3 methylation in these processes (**Supplementary Fig. 7**).

**Figure 2:**
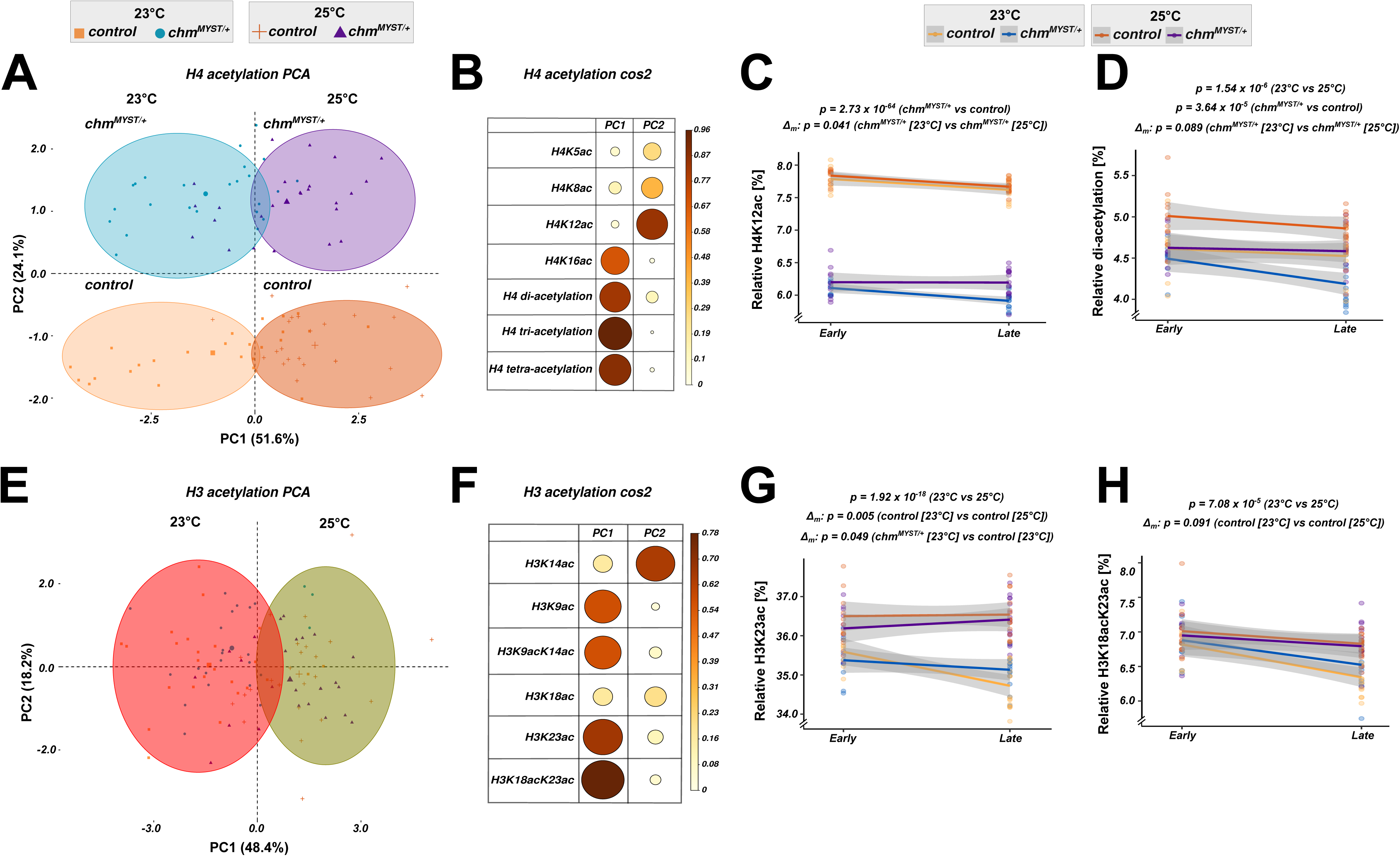
Histone PTM profiling indicates that temperature and starvation-specific changes in H3 and H4 acetylation. A) PCA of H4 acetylation data from mass spectrometry between control and *chm^MYST/+^*male mutants at 23°C and 25°C in early (0-10 hrs) and late (24-48 hrs) time points (N = 5, unpaired) B) cos2 plot of A) with PC1 and PC2 contributions for H4 acetylation C) H4K12ac and D) H4 di-acetylation dynamics between control and *chm^MYST/+^*male mutants at 23°C and 25°C in early (0-10 hrs) and late (24-48 hrs) time points (N = 5). E) PCA of H3 acetylation data from mass spectrometry between control and *chm^MYST/+^*male mutants at 23°C and 25°C in early (0-10 hrs) and late (24-48 hrs) time points (N = 5, unpaired) F) cos2 plot of E) with PC1 and PC2 contributions for H3 acetylation G) H3K23ac and H) H3K18acK23ac dynamics between control and *chm^MYST/+^*male mutants at 23°C and 25°C in early (0-10 hrs) and late (24-48 hrs) time points (N = 5, unpaired) *Non-significant values are not shown and significant p-values are mentioned in the figure. All statistical tests were obtained from linear regression model between relative percentage and condition (genotype_temperature) with time as an interaction term*

These observations suggest that histone acetylation, rather than methylation serves as a metabolic sensor in flies (**Charidemou & Kirmizis, 2024**). Collectively, our detailed profiling of histone PTM indicates that a temperature increase of 2°C is sufficient to at least partially rescue the impaired histone acetylation in fruit flies in a genotype specific manner.

### Transcriptomic data shows temperature- and genotype-specific differential expression

As *chm^MYST/+^* flies show a different transcriptional response to starvation at 23°C, we wondered whether this is also observed at 25°C (**Supplementary Fig. 8A**). Similar to the response seen at 23°C, we also observed a distinct clustering in PCA into 4 groups with PC1 contributing to nutrient status, while PC2 to genotype at 25°C (**Supplementary Fig. 8B; Venkatasubramani et al., 2023**). A gene-set enrichment analysis (GSEA) of the transcriptomic data revealed a similar transcriptional response to starvation in control and *chm^MYST/+^* male flies at 25°C compared to what we have seen at 23°C (**Supplementary Fig. 8C-D, Dataset 2-3**). However, when we looked at the starvation induced expression changes of individual genes at different temperatures, we observed only a poor correlation (Spearman r = 0.30 and 0.34 for control and *chm^MYST/+^* respectively) (**Fig. 3A**). With regards to the significant genes, we observed a number of genes that were uniquely regulated by starvation in a given genotype or temperature genes and a similar number of genes that were commonly regulated (**Fig. 3B**). Over-representation analysis (ORA) of shared significant genes from both genotypes mostly identified genes involved in metabolic processes (**Supplementary Fig. 9A-B, Dataset 4-5**).

**Figure 3:**
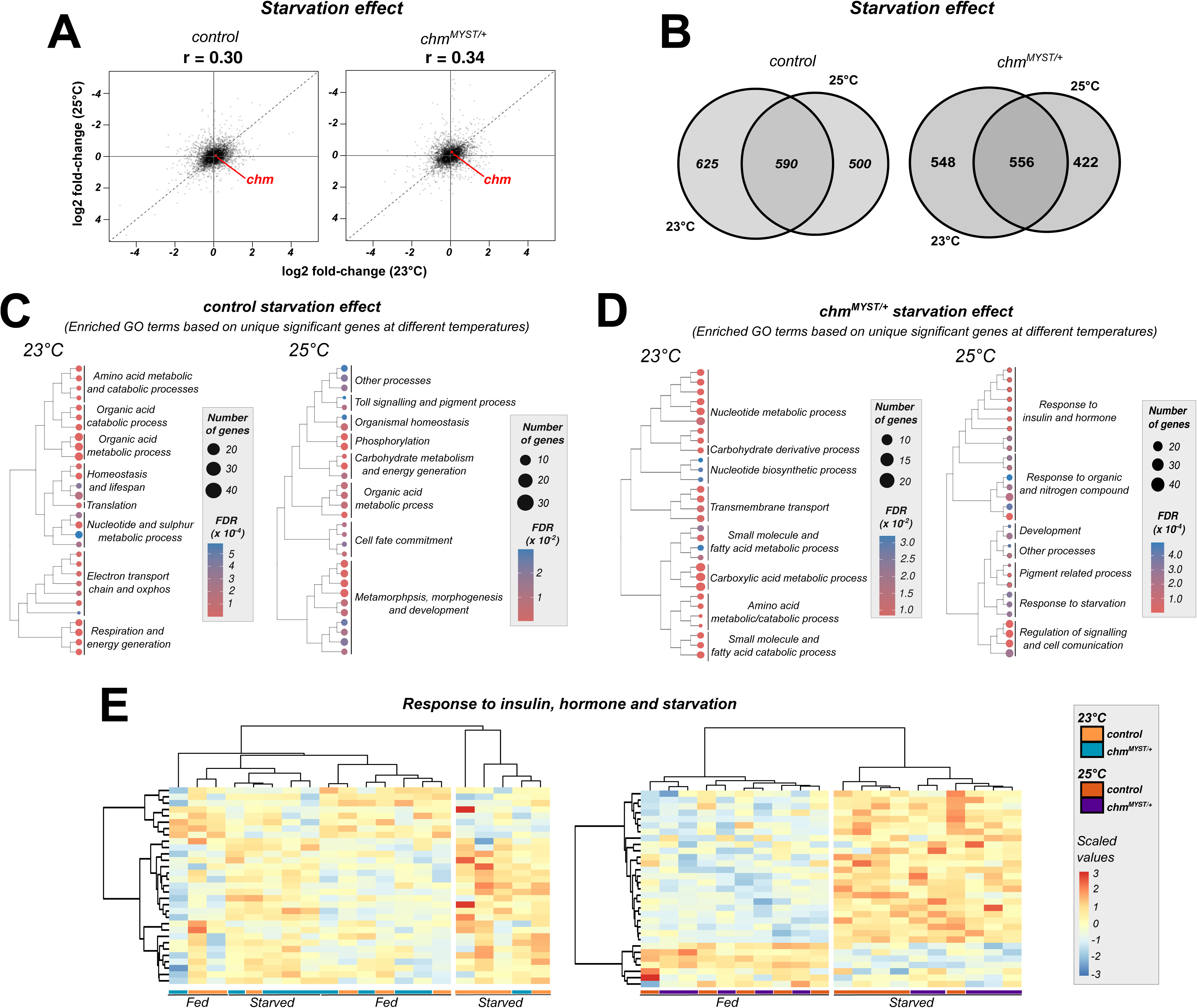
Transcriptomic data shows temperature-genotypic specific differences in genes related to insulin and hormonal response. A) log2FC-log2FC correlation plot of control and *chm^MYST/+^* starvation at 23°C (data obtained from **Venkatasubramani et al., 2023**) and 25°C (N = 5, unpaired) B) Venn diagram showing the number of unique and shared significant genes (FDR < 0.05) upon starvation in control and *chm^MYST/+^* starvation at 23°C and 25°C C-D) Treeplot showing the top 30 siginificant GO terms (FDR < 0.05) from over-representation analysis of unique significant genes in control and *chm^MYST/+^*starvation at 23°C (left) and 25°C (right) respectively (GO terms were clustered based on semantic similarity and the terms that were represented the most within a cluster was mentioned) E) Heatmap of scaled read counts of annotated genes from insulin, starvation and hormonal response in control and *chm^MYST/+^* starvation at 23°C (left) and 25°C (right)

The genes that were only differentially regulated in controls in a temperature dependent manner identified genes encoding for proteins involved in carbohydrate metabolism at 25°C, while the genes regulated at 23°C were enriched for mitochondrial processes (**Fig. 3C, Dataset 6-7**). In *chm^MYST/+^*flies, genes involved in insulin, hormone and starvation response were differentially regulated at 25°C, while an ORA revealed an enrichment of metabolic processes in genes differentially regulated at 23°C (**Fig. 3D, Dataset 8-9**). Interestingly, numerous studies have implicated insulin and other hormonal signaling in temperature control, proper feeding and stress resistance. Insulin secretion has also been liked to neuronal circuit which senses temperature fluctuation in *Drosophila* (**Enell et al., 2010; Koyama et al., 2020; Rauschenbach et al., 2014; Sudhakar et al., 2020**). We therefore further assessed the genes from insulin, hormone and starvation response. This led to a clustering of the samples with regard to their nutrient status at 25°C, while at 23°C, gene expression changes upon starvation largely clustered according to genotype (**Fig. 3E**). These results suggest that Chm has a much stronger effect on gene expression at 23°C than at 25°C, which is also the case for the histone acetylation profiles. As changes in these genes most likely results in differences in metabolism, we therefore decided to assess the changes in carbohydrates and lipids.

### Weight loss and trehalose induction upon starvation shows temperature- and genotype-specific differences

When fed ad libitum *chm^MYST/+^*flies are lighter when compared to corresponding control flies at 23°C (**Venkatasubramani et al., 2023**). Interestingly, this reduction in weight was no longer observed at 25°C (**Fig. 4A**). Furthermore, whereas *chm* mutants show a smaller weight loss upon starvation than wildtype flies at 23°C, the weight loss was similar between the genotypes at 25°C (**Fig. 4B**). This suggests that Chm is not strictly involved for nutrient homeostasis and the breakdown of macro molecules at 25°C.

**Figure 4:**
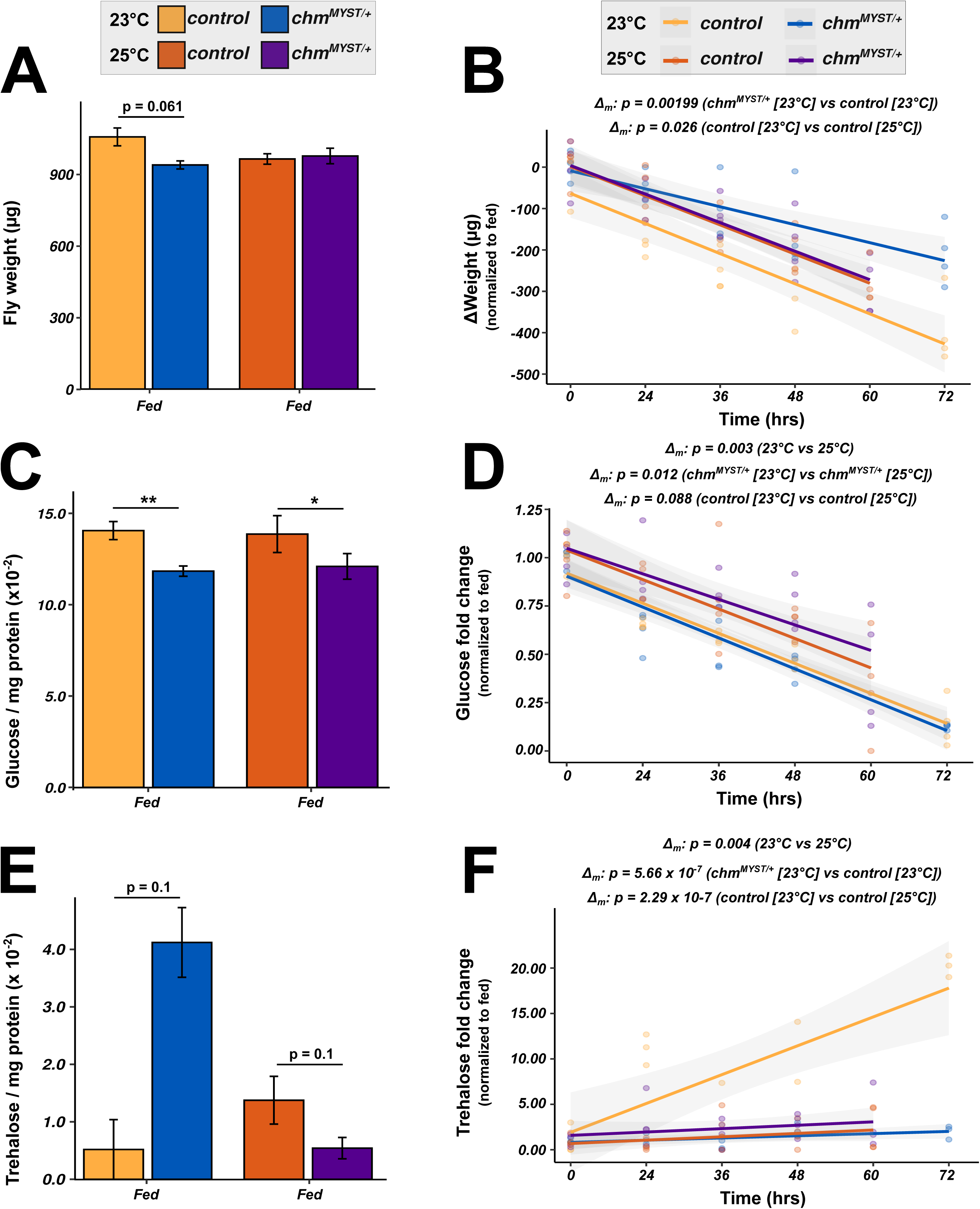
chm mutant flies showed impaired weight and trehalose homeostasis upon starvation at 23°C. A, Weight (per fly) C) Glucose (per mg protein), E) Trehalose (per mg protein) in control and *chm^MYST/+^* flies at 23°C and 25°C under ad libitum conditions respectively (N = 4 and 3 (for 23°C Trehalose), paired, at least 30 flies per replicate) B) Change in weight, D) Glucose fold change and E) Trehalose fold change upon starvation in control and *chm^MYST/+^* flies at 23°C and 25°C. All values are normalized to the fed condition of corresponding genotype and temperature (N = 4 and 3 (for 23°C Trehalose), paired, at least 30 flies per replicate) *Non-significant values are not shown. Paired t-test with FDR correction was performed for A), C) and E), while for B), D) and F), statistical tests from linear regression model between weight difference and condition (genotype_temperature) or weight difference and temperature were performed correspondingly. Error bars represent SEM*

Temperature differences have been shown to affect the synthesis and degradation of carbohydrates such as glucose, trehalose, glycogen and triacylglycerides (TAGs) (**Klepsatel et al., 2019).** We therefore measured these molecules in control and *chm^MYST/+^* flies under fed and different time-points of starvation and at different temperatures. It is noteworthy that *chm^MYST/+^* flies showed reduced levels of glucose irrespective of the temperature (**Fig. 4C**), which might explain the improved longevity of *chm^MYST/+^* at both temperatures (**Venkatasubramani et al., 2023**), as similarly low glucose levels are observed under caloric restriction. The reduction in glucose upon starvation were primarily affected by temperature rather than genotype (**Fig. 4D**) In contrast to glucose, the level of storage macro-molecules like TAGs and glycogen, were unaffected by both genotype or temperature but, as expected, showed a decreasing trend upon starvation (**Supplementary Fig. 10A-D**).

Unlike mammals, where glucose is the primary circulating sugar, fruit flies and nematodes contain a specific disaccharide called trehalose as the most abundant circulating sugar. Interestingly, under fed conditions, *chm^MYST/+^*flies showed higher levels of trehalose at 23°C but not at 25°C (**Fig. 4E**). Upon starvation, trehalose was induced in wildtype flies but not in *chm^MYST/+^*flies, most likely due to its already high levels under fed condition. At 25°C, in contrast, both control and *chm^MYST/+^* showed a similar induction of trehalose, albeit not as strong as control flies at 23°C (**Fig. 4F**).

These results prompted us to assess the RNA expression profiles of genes from trehalose metabolism in control and *chm^MYST/+^* flies at both temperatures. However, we failed to observe specific genes that were affected by the loss of *chm* and temperature (**Supplementary** Fig. 11), and speculate that the observed molecular changes could be a function of various small changes within a process rather than specific gene expression differences. Coincidentally, we also checked the proteome and acetylome data of control and *chm^RNAi^* at 23°C from our earlier study (**Venkatasubramani et al., 2023**). Intriguingly, we identified many proteins from trehalose metabolism that were differentially regulated upon the loss of Chm, such as GlyP, Pepck1, Hex-A, Tps1, UGP and Treh (**Supplementary Fig. 12**). Moreover, acetylation of GlyP, Pepck1, Tps1 and UGP, were reduced or completely lost in *chm* mutants (**Supplementary Fig. 13**). While, we do not have any information on the PTM status of these proteins at 25°C or their molecular function in starvation/temperature response, we postulate that increased metabolism and acetyl-CoA availability might aid in differential acetylation of and by Chm in these conditions.

### Overexpression of chm rescues trehalose induction and weight loss phenotype upon starvation at 23°C

In order to validate the observed phenotypes, we ectopically expressed *chm* in *chm^MYST/+^* background using *da-Gal4*. We have already shown that ectopic expression increases weight of flies at *ad libitum* and improves survival during starvation at 23°C (**Venkatasubramani et al., 2023**). Although not statistically significant, ubiquitous ectopic expression of *chm* marginally improved the weight loss upon starvation in *chm^MYST/+^*background (**Fig. 5A**). Furthermore, the ectopic expression also rescued the trehalose levels at fed and the inability of *chm* mutants to induce trehalose upon starvation (**Fig. 5B-C**) in a manner similar to the control flies as observed before. Glucose showed a decreasing trend upon starvation irrespective of the genotype, however, ectopic *chm* expression did not alter the levels of glucose at fed condition (**Fig. 5D-E**). In summary, these results highlight and validate the role of Chm in regulating weight and trehalose levels at lower temperatures.

**Figure 5:**
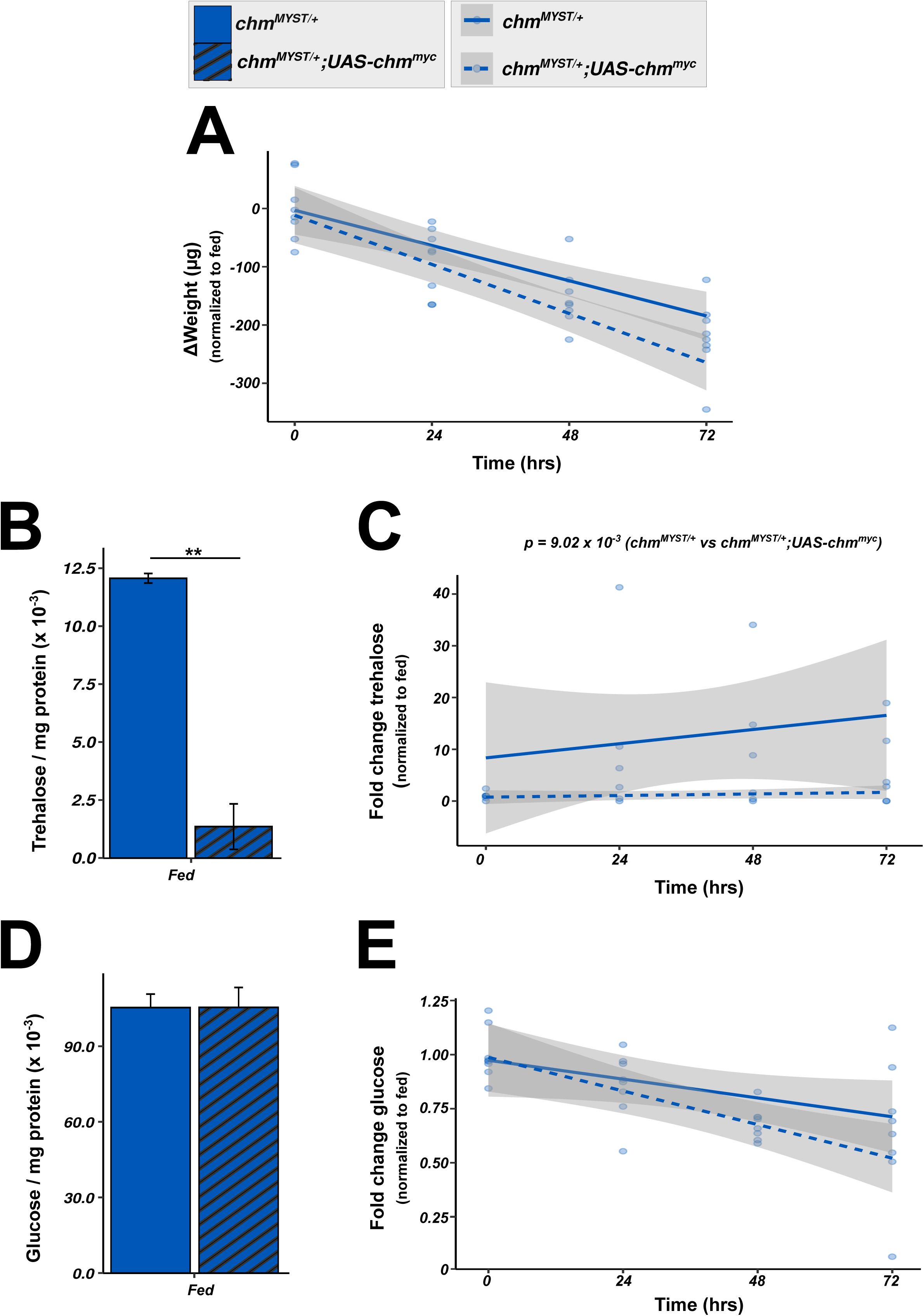
Weight and trehalose homeostasis is improved upon chm overexpression at 23°C. A) Change in weight, C) Trehalose fold change and E) Glucose fold change upon starvation in *chm^MYST/+^* and *chm^MYST/+^;UAS-chm^myc^*flies at 23°C. All values are normalized to the fed condition of corresponding genotype and temperature (N = 4 and 3 (for Trehalose), paired, at least 30 flies per replicate) B) Trehalose and D) Glucose in *chm^MYST/+^* and *chm^MYST/+^;UAS-chm^myc^*flies at 23°C under ad libitum conditions respectively (N = 4 and 3 (for 23°C Trehalose), paired, least 30 flies per replicate) *Non-significant values are not shown. Paired t-test was performed for B) and D), while for A), C) and E), statistical tests from linear regression model between weight difference and condition (genotype) were performed. Error bars represent SEM*.

## Discussion

In this study, we showed that Chm mediates starvation resilience only at lower temperatures and does not play a role at higher temperatures. At 23°C *chm* mutants show a change in H4 acetylation, an altered expression of genes linked to insulin and starvation response, a lower weight and a dampened up-regulation of trehalose upon starvation. Most alterations are not manifested at higher temperatures, suggesting that Chm’s function in adult flies is restricted to defined environmental conditions.

Our comprehensive histone analysis suggested a potential role of histone acetylation in regulating the response to starvation at different temperatures. Specifically, *chm^MYST/+^*flies at lower temperatures showed differing trends of H4K12ac and H4-diacetylation upon starvation. In fact, these changes were observed to a stronger extent when comparing middle-aged fed wild-type flies at 18°C and 25°C (**Peleg et al., 2016**), further validating the influence of temperature on histone acetylation, especially at H4.

Transcriptomic analysis showed that wildtype flies respond surprisingly different to starvation when raised at 23°C (**See Fig. 5D and 5F in Venkatasubramani et al., 2023**) or 25°C (**Supplementary Fig. S14A-B**). While *chm* mutants have an effect on transcription under both temperatures, the genes involved in the response to hormone, insulin and starvation were affected by a *chm* mutation at 23°C but not at 25°C. The same is true for the up-regulation of trehalose upon starvation, which we observed at 23°C but not at 25°C. In fact, similar to our previous studies, which showed that *chm* mutant flies are already stressed under well fed conditions, the *chm^MYST/+^* flies have an elevated trehalose level at 23°C. This goes well in line with previous studies that indicate the accumulation of trehalose upon stress in various organisms from yeast to plants and invertebrates. Furthermore, trehalose homeostasis is controlled by hormones depending on the environmental conditions. ILPs aid in trehalose homeostasis by controlling the expression of Treh or by controlling its activity by binding to the enzyme. In addition, other hormones such as Juvenile and ecdysone hormone also affects the expression of Treh and Tret-1 (**Bandara et al., 2009; Bobrovskikh & Gruntenko, 2023; Jang & Lee, 2018; Kosar et al., 2019; Santos et al., 2024; Tellis et al., 2023; Thorat et al., 2012; Xu et al., 2013**). These observations align well with the observed transcriptional and molecular changes at 25°C of *chm^MYST/+^*mutants.

### What can cause the temperature specific changes in histone acetylation levels?

Temperature changes have been shown to affect the activity of enzymes, including metabolic enzymes. Such changes in biochemical properties could occur via moderation of substrate concentration or via post-translational modifications of the corresponding proteins (**Johnston & Dunn, 1987; Peleg et al., 2016; Yamauchi et al., 1975**). It is possible that at lower temperatures, the metabolic rates are lower, thus the flux that maintains acetyl-CoA, the precursor for acetylation reactions, is lower. This could alter the capacity of Chm to maintain the homeostatic levels of specific histone acetylation during starvation or at times of stress. It is noteworthy to mention that HBO1, the orthologue of Chm, has a lower affinity (high kM) to acetyl-CoA as compared to other histone acetyltransferases such as HAT1, MOF, Tip60 and p300/CBP (**Ronowska et al., 2018; Tan et al., 2022; Wu et al., 2012**). Following up and characterizing the acetyl-CoA cellular flux will further assess whether lower temperature-dependent metabolic fluxes, combined with lower Chm are the culprit behind the impaired ability of *chm* mutants to survive starvation at lower temperatures.

In conclusion, our data shows the role of acetyltransferase Chm in fine tuning response to starvation at a resolution of 2°C. Its important role in withstanding periods of starvation at colder times may underlie the natural selection that led to its current expression levels in flies, although flies with lower expression are long-lived when food is abundant. In fact, it is possible that flies from different population might have differential *chm* expression due to the presence of different significant SNPs in *chm* gene region. As predicted by these studies, this work reveals the importance and role of *chm in* novel environments (**Croze et al., 2017; Levine & Begun, 2008**). Further studies could explore the function and relevance of Chm in stress and metabolic regulations from fly populations of varying environmental conditions.

## Materials and Methods

### Fly maintenance

Fly lines were maintained in a incubator (Panasonic, MLR-235H-PE) at 25°C with a 12 hr/12 hr light-dark cycle at 60% relative humidity. Composition of the fly food and comparison between Germany and Japan is given in **Table.1.** Details of fly lines used in the study are given in **Table.2**.

**Table. 1:** Composition of fly food (Munich)

| Components | Amount (Germany) | Amount (Japan) |
| --- | --- | --- |
| Agarose | 130.00 g | 187.20 g |
| Corn meal | 1300.00 g | 1600.00 g |
| Soy flour | 150.00 g | 160.00 g |
| Yeast | 300.00 g | 296.00 g |
| Maltase | 650.00 g | 640.00 g |
| Molasses | 1300.00 g | 640.00 g |
| Methyl 4-Hydroxybenzoate (Nipagin) | 415.00 ml Ethanol (10%) | 320.00 ml Ethanol (10%) |
| Acid mix | 295.00 ml | - |
| 85% phosphate | - | 19.20 ml |
| Water | Up to 16 l | Up to 16 l |

**Table. 2:**
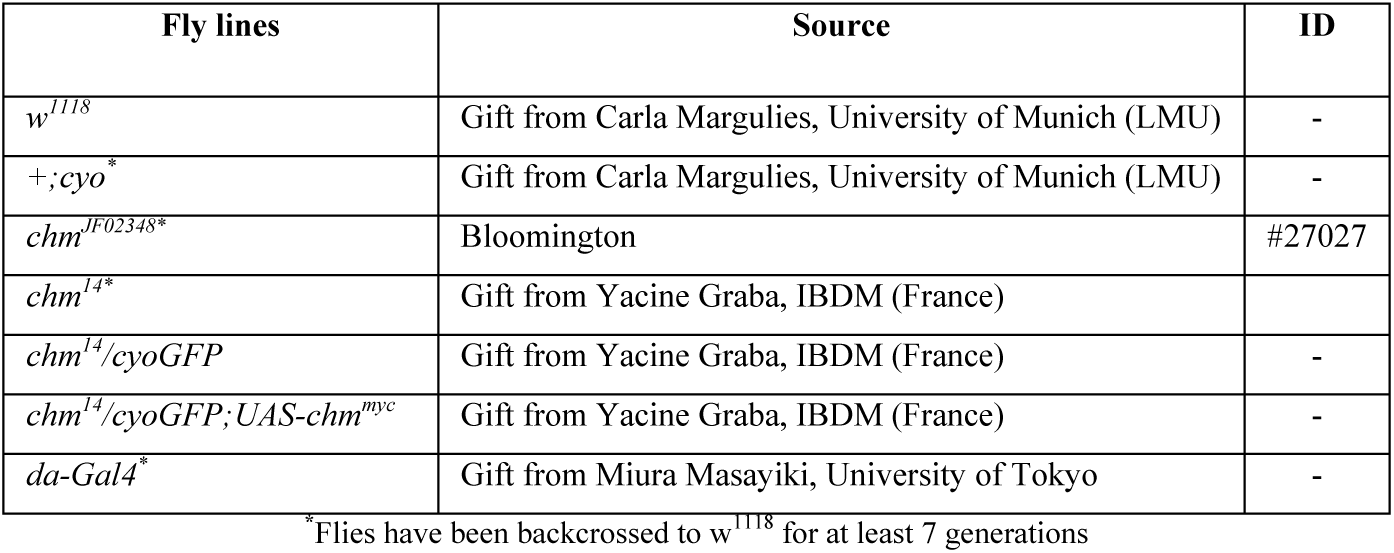
Details of fly lines used in this experiment.

### Starvation assay

Flies of age 8-9 days old developed, raised and starved at corresponding temperatures with 60% relative humidity and 12 hr light/12 hr dark cycle were used for starvation experiments. Flies (30 flies - small vial; 50 flies - big vial) were transferred to an empty fly vial containing a tissue soaked in water (1 ml - small vial; 4 ml - big vial). Total number of flies for each experiment ranged from 100-200 for each genotype/condition tested. Readings were taken until flies were dead in all genotypes with dead flies counted every 10-12 hr. Experiments at 23°C and 25°C (in males) were performed in both Japan and Germany to exclude any differences in food and other laboratory conditions. For females, starvation at 23°C and 25°C were performed only in Germany. Furthermore, starvation at 18°C, 21°C and 27°C in both males and females were performed only in Japan.

### Weight measurement

Flies of the fed and starved conditions were transferred to a 2.0 ml tube and snap-frozen in liquid nitrogen. Following this, weight of each empty tube was measured. Heads and bodies were obtained by passing through sieves. First sieve (width: 710 um) separates the bodies from remaining and the second (width: 355 um) separates heads from wings/legs (Analysensieb). 20-30 heads and bodies were transferred to the corresponding 1.5 ml tube and the weight was measured again. Difference in weight between the two was considered as the corresponding weight of 20-30 flies from which weight per fly was calculated. Weights were measured using KERN ABJ 120-4NM weighing machine. Samples and all the components were kept in dry ice for the entire duration of the experiment.

### Post-translational modification of histones

#### Sample preparation

Fly heads were obtained following the same procedure given in *Body and head weight measurement* section. Approx. 300 μl of homogenization buffer [60 mM KCl, 15 mM NaCl, 4 mM MgCl_2_, 15 mM HEPES (pH 7.5), 0.5% Triton-X-100, 0.5 mM DTT, 20 mM sodium butyrate and 1 tablet protease inhibitor] were added to 30-50 fly heads and were homogenised extensively with an electrical stirrer (5 x 10 s ON and 15 s OFF). Following this, sonication was performed with Bioruptor^®^ Pico for 3 x 10 s ON and 45 s OFF at 4°C. Obtained lysate was centrifuged at 20,817 x g for 30 min. Obtained pellet was resuspended in 200 μl of 0.2 M H_2_SO_4_, vortexed heavily and rotated overnight at 15 rpm and 4°C. Subsequently, overnight incubated lysate was centrifuged at 20,817 x g (max. Speed) for 10 min at 4°C. Histone were precipitated by adding trichloroacetic acid (TCA) (ThermoScientific, Cat. No 85183) to reach 26% final concentration. Tubes were mixed and incubated at 4°C for 2 hr and spun at 20,817 g for 15 min. Pellets were washed thrice with ice-cold 100% acetone (VWR, Cat. No AA22928-K2) (5 min rotation at 4°C, 15 min of 20,817 g spin at 4°C between washes), dried for 15 min at room temperature and resuspended in 20 μl of 1x Laemmli sample buffer for million cell and boiled at 95°C for 5 min. Samples were stored at -20°C until further use. The histones corresponding to 0.5 million cells were separated onto 4–20% pre-cast polyacrylamide gels (SERVA, Cat. No 43277.01). Gels were briefly stained with InstantBlue Coomassie Protein Stain (abcam, Cat. No ab119211). For targeted mass-spectrometry analysis, histones bands were excised, washed once with MS-grade water (Sigma-Aldrich, Cat. No 1153331000) and de-stained twice (or until transparent) by incubating 30 min at 37°C with 200 μl of 50% acetonitrile (ACN) (Carl Roth, Cat. No 8825.2) in 50 mM Ammonium bicarbonate (NH_4_HCO_3_) (Carl Roth, Cat. No T871.1). Gel pieces were then washed twice with 200 μl MS-grade and twice with 200 μl of 100% ACN to dehydrate them. Histones were in-gel acylated by first adding 20 μl of d6 acetic anhydride (Sigma-Aldrich, 175641-5G), followed by 40 μl of 100 mM NH_4_HCO_3_. After 5 min, 140 μl of 1 M NH_4_HCO_3_ was slowly added to the reaction. pH of the final solution should be around 7 (In cases where pH was acidic, few microlitres of 1 M NH_4_HCO_3_ was added). Samples were incubated at 37°C for 45 min at 550 rpm. Following this, samples were washed 5 times with 200 μl of 100 mM NH_4_HCO_3_, 4 times with 200 μl of MS-grade water and 4 times with 200 μl of 100% ACN. They were spun down briefly and all remaining ACN was removed. Gel pieces were rehydrated in 50 μl of trypsin solution (25 ng/ mL trypsin in 100 mM NH_4_HCO_3_) (Promega, Cat. No V5111) and incubated at 4°C for 30 min. After the addition of 150 μl of 50 mM NH_4_HCO_3_, histones were in-gel digested overnight at 37°C at 550 rpm. Peptides were sequentially extracted by incubating 10 min at room temperature with 150 μl of 50 mM NH_4_HCO_3_, twice with 150 μl of 50% ACN (in MS-grade water) 0.1% trifluoroacetic acid (TFA) and twice 100 μl of 100% ACN. During each of the above washing step, samples were sonicated for 3 min in a water bath followed by a brief spin down. Obtained peptides were dried using a centrifugal evaporator and stored at -20°C until resuspension in 30 μl of 0.1% TFA. For desalting, peptides were loaded in a C18 Stagetip (prewashed with 20 μl of methanol followed by 20 μl 80% ACN 0.1% TFA and equilibrated with 20 μl of 0.1% TFA), washed 2 times with 20 μl of 0.1% TFA and peptides were eluted 3 times with 10 μl of 80% ACN 0.25% TFA. Flow through obtained from loading of peptides in C18, were further desalted with TopTip Carbon (glygen, Cat, No TT1CAR.96) by loading the flow through thrice (prewashed thrice with 30 μl of 100% ACN followed by equilibration thrice with 30 μl of 0.1% TFA), washed 5 times with 30 μl of 0.1% TFA and eluted thrice with 15 μl of 70% ACN and 0.1% TFA. Eluted peptides from both desalting steps were combined and evaporated in a centrifugal evaporator, resuspended in 15-17 μl of 0.1% TFA and stored at -20°C until mass spectrometry analysis.

#### Targeted mass spectrometry

Desalted histone peptides in 0.1% TFA were injected in an RSLCnano system (Thermo Fisher Scientific) and separated in a 15-cm analytical column (75μm ID home-packed with ReproSil-Pur C18-AQ 2.4 μm from Dr. Maisch) with a 50-min gradient from 4 to 40% ACN in 0.1% formic acid at 300 nl/min flowrate. The effluent from the HPLC was electrosprayed into Q Exactive HF mass spectrometer (Thermo Fisher Scientific). The MS instrument was programmed to target several ions except for the MS3 fragmentation (22). Survey full scan MS spectra (from m/z 270-730) were acquired with resolution R=60,000 at m/z 400 (AGC target of 3x106). Targeted ions were isolated with an isolation window of 0.7 m/z to a target value of 2x105 and fragmented at 27% normalized collision energy. Typical mass spectrometric conditions were: spray voltage, 1.5 kV; no sheath and auxiliary gas flow; heated capillary temperature, 250°C.

#### Data analysis

Raw data from mass spectrometry was analysed using Skyline (**Pino et al., 2020**) v21.1. Peak integration was performed for H3 and H4 peptides for each of its corresponding modifications. Relative levels of each PTM were calculated from the obtained intensities using R environment, based on the formula given in (**Feller et al., 2015**).

### RNA-sequencing

#### RNA extraction

30 heads were homogenized with an electrical stirrer with 500 μl of Trizol (Thermo Fisher; cat. no. 15596026). Chloroform was added at the ratio of 1:5 with Trizol and the solutions were mixed for 15 s by inverting the tubes. After 5 min incubation at room temperature, samples were centrifuged at 12,000 x g for 15 min. Aqueous phase was transferred to a new tube from the centrifuged sample to which isopropanol was added at 1:1 ratio of the obtained aqueous phase, vortexed briefly and incubated for 10 min at room temperature. They were then centrifuged for 10 min at 12,000 x g. Supernatant was discarded and obtained pellet was washed with 750 μl of 80% ethanol. After brief vortexing, samples were centrifuged at 8,000 x g for 5 min. Obtained supernatant was discarded and pellet was air dried for 5 min inside the hood and resuspended in RNase-free water.

#### Library preparation

1 ug of RNA, obtained from fly heads was used for library preparation. Both total and mRNA quality was assessed on a 2100 Bioanalyzer (Agilent Technologies, Cat. No G2939BA) using RNA pico assay kit (Agilent RNA 6000 Pico Kit, cat. No: 5067-1513) using manufacturer’s protocol. rRNA depletion was performed using NEBNext rRNA Depletion Kit (Human/Mouse/Rat) (NEB #E6310) and library preparation for RNA-sequencing was performed using NEBNext Ultra II Directional RNA Library Prep Kit for Illumina (NEB #E7760) following manufacturer’s protocol. Libraries were sequenced on an Illumina HiSeq 1500 instrument at the Laboratory of Functional Genomic Analysis (LAFUGA, Gene Center Munich, LMU).

#### Data analysis

A total of 50 bp paired-end reads were aligned to the D. melanogaster reference genome (release 6) using STAR aligner (version 2.5.3a) with providing GTF annotation (dmel-all-r6.17.gtf). Reads with multiple alignments were filtered by setting outFilterMultimapNmax parameter to 1. Reads were counted per gene with parameter –quantMode GeneCounts. BAM files were converted to normalized bedgraph coverages using genomeCoverageBed command (bedtools version 2.27.1) with -scale parameter set to divide by the total number of reads and multiplied by a million. Bedgraph files were converted to tdf files (igvtools version 2.3.98) to visualize in the IGV browser.

Count tables (read counts per gene) were read into R environment and low count genes were filtered out (at least 3 read per gene in 10% of the samples analyzed together). Differential expression analysis was performed by DESeq2 (**Love et al., 2014**) package (version 1.24) by adding replicate information as batch variable. Samples that were directly compared to each other were fitted in the same DESeq2 model. Log2FoldChange estimates and adjusted P-values were obtained by the results function (DESeq2) and an FDR cutoff < 0.05 was applied. In addition, the less stringent Π-value that includes both statistical and biological information was used. Π-value takes into consideration log2FoldChange and p-value to obtain values between 0 and 1 (**Hostrup et al., 2022; Xiao et al., 2014**). For principal component analysis (PCA) analysis batch effect was corrected by the remove batch effects function from limma (**Ritchie et al., 2015**) (package version 3.52.0) on the normalized read counts.

Gene Set Enrichment Analysis was performed on the obtained results from different conditions/comparisons using the gseGO function from clusterProfiler (**Yu et al., 2012**) (package version 3.12.0) by ranking the genes based on t-statistic value without any log2FoldChange or p-adjusted cut-off. GSEA plots with selected GO terms were also generated with R environment with these selected GO terms having FDR < 0.05 cut-off.

### Carbohydrate (Glucose, Glycogen and Trehalose) and TAG quantification

This methodology was adopted and modified from (**Tennessen et al., 2014**), with minor modifications. Briefly, 10-15 flies were snap-frozen in liquid nitrogen and lysed with 1x PBS (for carbohydrate) and 1x PBST-0.05% (for TAGs) using an electric stirrer. An aliquot from the obtained lysate was used for protein quantification using Quick Start™ Bradford 1x Dye Reagent (BIO-RAD, cat no: 5000205). Following heat treatment, the lysate was stored at -80°C until analysis.

For glycogen and trehalose, samples were treated either with amyloglucosidase (Sigma-Aldrich, cat no: 11202332001) or with porcine trehalase (Sigma-Aldrich, cat no: T8778-1UN). In addition, for both glycogen and trehalose, non-enzyme treated control, to measure the basal glucose, was also used by replacing the corresponding enzyme with 1x PBS or 1x Trehalase Buffer (5 mM Tris pH 6.6, 137 mM NaCl, 2.7 mM KCl). Accordingly, glucose, glycogen and trehalose standards were prepared from 0 to 0.16 mg/ml range. For glycogen, samples were diluted between 1:3 or 1:5 and the enzymatic treatment was carried out for 60 minutes at 37°C, while for trehalose, samples were either undiluted or diluted 1:2 and the enzymatic treatment was carried out overnight at 37°C. Following this, absorbance at 340 nm was measured using a plate reader. Amount of glycogen and trehalose was calculated by measuring the difference between enzyme treated and non-treated samples and normalized to protein amounts. In cases where the absorbance fall below the standard curve, the value was replaced with zero as they were below the threshold limit of the quantification method.

For TAGs, samples were treated with triglyceride reagent (Sigma-Aldrich, cat no: T2449). In addition, non-enzyme treated control, to measure the basal glycerol, was also used by replacing the corresponding enzyme with 1x PBST. Accordingly, glycerol standards were prepared from 0 to 1.0 mg/ml range using 2.5 mg/ml triolein equivalent glycerol standard solution (Sigma-aldrich, cat no: G7793). For quantification of TAGs, samples were undiluted and the enzymatic treatment was carried out for 60 minutes at 37°C. Following this, absorbance at 540 nm was measured using a plate reader. Amount of TAG was calculated by measuring the difference between enzyme treated and non-treated samples and normalized to protein amounts. In cases where the absorbance fall below the standard curve, the value was replaced with zero as they were below the threshold limit of the quantification method.

### Food consumption

#### Preparation of agar based fly food with 0.5% brilliant blue FCF

This methodology was adopted and modified from (**Shell et al., 2018**). The recipe shown in **Table. 3** is for 1 l of the agar-based fly food. Mix agar and distilled water by heating and stirring (use magnetic stirrer and heater) them continuously until agar is dissolved.

**Table. 3:** Composition of fly food for food consumption assay.

| Components | For 1 litre |
| --- | --- |
| Water | 640 ml |
| Agar | 29 g |
| Apple juice | 280 ml |
| Rubensirup | 40 ml |
| 10% Nipagin (in ethanol) | 22.5 ml |
| Brilliant FCF (0.5% w/v) | 5g |

Once agar is dissolved, apple juice and Rubensirup was added and mixed well. Following this, brilliant blue FCF was also added and all these components were mixed until dissolved. Once the temperature reaches around 70°C, Nipagin was added little by little with constant stirring. Approximately 9 ml of the fly food was added to a 60 mm petri dish and kept at room temperature (around 25°C) until solidified. The petri dishes were then stored at 4°C until use. Before transferring the flies for experiment, the food was kept to room temperature for at least 1 hour.

Measurement of consumed and excreted food: Once the food is warm, 15-20 flies were transferred to an empty fly vial and the open end was sealed with the petri-dish containing fly food. Cellotape was used to make sure that the petri dish doesn’t get disturbed. The flies were then kept at 23°C, with a 12 hr - 12 hr day-light cycles and 60% humidity for 24 hr. Flies were then collected and transfered to an eppendorf. 200 μl of dH2O were added and homogeneized using electrical homogeneizer, followed by centrifugation at full speed for 5 minutes. Supernatant was transferred and the insoluble material was discarded. Obtained supernatant was made up the volume to 1.5 ml with dH2O. To measure the amount of food excreted, 5 ml of dH2O was added the sides of the fly vial and pipetted up and down with a 1 ml pipette to make sure all the colored excreta were resuspended and the obtained solution was transferred to an eppendorf. Both intake and excreted food absorbance was measured at 630 nm in a plate reader with dH2O as blank. To measure total food consumed, the absorbance from intake and excreted food was added.

### Plots and statistical analysis

All statistical analysis was performed in R (R Core Team (2021). R: A language and environment for statistical computing. R Foundation for Statistical Computing, Vienna, Austria. URL https://www.R-project.org/) environment unless otherwise mentioned. Graphics in all figures were created using Biorender.com. Statistical tests were decided based on the experimental design and measurements and have been mentioned in each of the figure legends. Plots and graphs generated for all experiments were also generated in R environment unless otherwise mentioned.

## Data availability

The datasets produced in this study are available in the following databases:

- *Transcriptomic data:* Gene Expression Omnibus GSE287187 (https://www.ncbi.nlm.nih.gov/geo/query/acc.cgi?acc=GSE287187)
- *Histone PTM mass spectrometry data:* ProteomeXchange Consortium via the PRIDE^20^ partner repository PXD035947 (http://www.ebi.ac.uk/pride/archive/projects/PXD035947)

## Acknowledgments

We would like to thank the members of the Imhof lab, Peleg lab, Tanimoto lab, Becker department, Andreas Ladurner and Raffaele Teperino for their inputs and suggestions, Catherine Regnard and Silke Krause from Becker department, Stefan Krebs and Helmut Blum from LAFUGA facility for sequencing, Nicolas Gompel and Christa Schwarzlose for fly food and maintenance respectively. We thank Tobias Straub, Tamas Schauer and Wasim Aftab for their assistance in experimental design, statistics and bioinformatic analysis. We thank Markus Hohle from QBM for his constant support.

AVV is supported by the QBM and SFB1309. TI was supported by the Ministry of Education, Culture, Sports, Science and Technology (MEXT): 21K06369. Work in the AI lab was funded by grants from the DFG, grant numbers 213249687 (CRC1064) and 325871075 (CRC1309). The Peleg lab is supported by the FBN, DFG grant (458246576), Longevity Impetus grant from Norn Group.

## Author contributions

AVV, SP and AI conceptualized the project. AVV discovered the starvation phenotype, conducted the experiments, performed data analysis and prepared all the figures. TI performed some of the starvation assays. IF performed LC/MS analysis for histone acetylation. HT provided technical expertise on starvation assays and temperature variation. AI and SP also supervised and AVV. AI, AVV and SP wrote the manuscript with comments from all authors.

## Supplementary figures

**Supplementary figure 1: Survival curves shows the trend of starvation in male control and chm^MYST/+^ flies at different temperature**

A-E) Kaplan-Meier plot of control and *chm^MYST/+^* male flies at 18°C (N = 3, paired), 21°C (N = 3, paired), 25°C (N = 5, paired) and 27°C (N = 3, paired). *Error bars represent SEM*.

**Supplementary** figure 2**: Survival curves shows the trend of starvation in female control and chm^MYST/+^ flies at different temperatures**

*A-*E) Kaplan-Meier plot of control and *chm^MYST/+^* female flies at 18°C (N = 3, paired), 21°C (N = 3, paired), 25°C (N = 4, paired) and 27°C (N = 3, paired). *Error bars represent SEM*.

**Supplementary figure 3:** *Pilot analysis suggests additive effect of temperature on starvation response in chm^MYST/+^ flies*

A) Experimental design for performing starvation assays at different combinations of 23°C and 25°C

B) Mean survival difference between control and *chm^MYST/+^* flies at different combinations of 23°C and 25°C (Data from 23°C were data obtained from **Venkatasubramani et al., 2023,** N = 5 [paired, for 25-25-25], N = 2 [paired, 25-25-23]) . *Error bars represent SEM*

**Supplementary figure 4:** *Survival curves shows the trend of starvation in control and chm^MYST/+^ flies at different temperatures in Japan*

Kaplan-Meier plot of control and *chm^MYST/+^*male flies at A) 23°C and B) 25°C (N = 1)

**Supplementary figure 5:** *Survival curves shows the trend of starvation in control and chm^RNAi^ flies at different temperatures*

Kaplan-Meier plot of control and *chm^RNAi^* A) male and B) female flies at 25°C (N = 4, paired). *Error bars represent SEM*

**Supplementary figure 6:** *Total food consumption is unaffected irrespective of temperature or genotype*

A) Food intake B) Excreted and C) Total consumed in control and *chm^MYST/+^* male flies at 23°C and 25°C respectively (N = 8, unpaired)

*Non-significant values are not shown. ANOVA followed by Tukey post-hoc test was performed, if ANOVA was significant. Error bars represent SEM*

**Supplementary figure 7:** *Histone methylation does shows any clustering based on the confounding factors*

Left) PCA of H3 methylation data from mass spectrometry between control and *chm^MYST/+^*male mutants at 23°C and 25°C in early (0-10 hrs) and late (24-48 hrs) time points (N = 5, unpaired), Right) cos2 plot of C) with PC1 and PC2 contributions for H4 methylation

**Supplementary figure 8:** *Transcriptome of 25°C starvation effect showed similarities to 23°C*

A) Experimental design for transcriptomic analysis of fed and starved control and *chm^MYST/+^* flies

B) PCA plot of control and *chm^MYST/+^* male flies at fed and starved condition and 25°C (N = 5, unpaired). Colors indicate the genotype and shapes indicate the nutrient status

C-D) Tree plot depicting the top 30 significant GO terms from GSEA of the starvation effect in control and *chm^MYST/+^*respectively. Color on the circles indicates enrichment and the size indicates number of genes annotated in that pathway (GO terms were clustered based on semantic similarity and the terms that were represented the most within a cluster was mentioned).

**Supplementary figure 9:** *Common significant genes from different temperatures showed enrichment of similar GO terms irrespective of the genotype*

A-B) Treeplot showing the top 30 significant GO terms (FDR < 0.05) from over-representation analysis of common significant genes between 23°C and 25°C in control and *chm^MYST/+^*starvation respectively (GO terms were clustered based on semantic similarity and the terms that were represented the most within a cluster was mentioned).

**Supplementary figure 10:** *TAG and glycogen is unaffected by temperature or genotype*

A) TAG (per mg protein) and C) Glycogen (per mg protein) in control and *chm^MYST/+^* flies at 23°C and 25°C under ad libitum conditions respectively (N = 4, paired, at least 30 flies per replicate)

B) TAG fold change and D) Glycogen fold change upon starvation in control and *chm^MYST/+^* flies at 23°C and 25°C. All values are normalized to the fed condition of corresponding genotype and temperature (N = 4, paired, at least 30 flies per replicate) *Non-significant values are not shown. Paired t-test with FDR correction was performed for A) and C), while for B) and D) statistical tests from linear regression model between weight difference and condition (genotype_temperature) or weight difference and temperature were performed correspondingly. Error bars represent SEM*

**Supplementary figure 11:** *Read counts of identified genes involved in trehalose metabolism from control and chm^MYST/+^ genotypes at 23*°C *and 25*°C *in fed (0 hr) and starved conditions (24 hr)* (N = 5, unpaired, Data for 23°C is obtained from **Venkatasubramani et al., 2023).** *Error bars represent SEM*

**Supplementary figure 12:** *LFQ of identified proteins in trehalose metabolism from control and chm^RNAi^ genotypes at 23°C in fed condition that are affected by loss of chm* (N = 4, paired, Data for 23°C is obtained from **Venkatasubramani et al., 2023).** *Error bars represent SEM*

**Supplementary figure 13:** *LFQ of identified acetyl sites from proteins in trehalose metabolism from control and chm^RNAi^ genotypes at 23°C in fed condition that are affected by loss of chm.* The acetylated lysine is marked in red color (N = 4, paired, Data for 23°C is obtained from **Venkatasubramani et al., 2023).** *Error bars represent SEM*

**Supplementary figure 14:** Heatmap of scaled read counts (right) and box-plot of variance-stabilization transformed read counts (left) of annotated genes from A) cellular response to stress and B) carbohydrate metabolic process in control and *chm^MYST/+^* starvation at 25°C respectively.

*Non-significant values are not shown. ANOVA followed by Tukey post-hoc test was performed, if ANOVA was significant*.

## Supplementary tables/dataset

**Dataset 1:** *Percentages of selected H3 and H4 post-translational modifications in control and chm^MYST/+^ flies at 23°C and 25°C and 0-48 hr starvation*

**Dataset 2:** *Significant GO terms (FDR < 0.05) from GSEA of transcriptome in control starvation at 25°C*

**Dataset 3:** *Significant GO terms (FDR < 0.05) from GSEA of transcriptome in chm^MYST/+^ starvation at 25°C*

**Dataset 4:** *Significant GO terms (FDR < 0.05) from over-representation analysis common genes in chm^MYST/+^ starvation at 23°C and 25°C*

**Dataset 5:** *Significant GO terms (FDR < 0.05) from over-representation analysis common genes in control starvation at 23°C and 25°C*

**Dataset 6:** *Significant GO terms (FDR < 0.05) from over-representation analysis unique genes in control starvation at 23°C*

**Dataset 7:** *Significant GO terms (FDR < 0.05) from over-representation analysis unique genes in control starvation at 25°C*

**Dataset 8:** *Significant GO terms (FDR < 0.05) from over-representation analysis unique genes in chm^MYST/+^starvation at 23°C*

**Dataset 9:** *Significant GO terms (FDR < 0.05) from over-representation analysis unique genes in chm^MYST/+^starvation at 25°C*

## References

Bandara, A., Fraser, S., Chambers, P. J., & Stanley, G. A. (2009). Trehalose promotes the survival of Saccharomyces cerevisiae during lethal ethanol stress, but does not influence growth under sublethal ethanol stress. FEMS Yeast Research, 9(8), 1208– 1216. 10.1111/j.1567-1364.2009.00569.x

Bhakti Chandegra, Jocelyn Lok Yee Tang, Haoyu Chi1, & Nazif Alic1. (2017). Sexually dimorphic effects of dietary sugar. Aging, 9(12), 2521–2528.

Bobrovskikh, M. A., & Gruntenko, N. E. (2023). Mechanisms of Neuroendocrine Stress Response in Drosophila and Its Effect on Carbohydrate and Lipid Metabolism. Insects, 14(5). 10.3390/insects14050474

Charidemou, E., & Kirmizis, A. (2024). A two-way relationship between histone acetylation and metabolism. Trends in Biochemical Sciences, 49(12), 1046–1062. 10.1016/j.tibs.2024.10.005

Chiang, M. H., Lin, Y. C., Chen, S. F., Lee, P. S., Fu, T. F., Wu, T., & Wu, C. L. (2023). Independent insulin signaling modulators govern hot avoidance under different feeding states. In PLoS Biology (Vol. 21, Issue 10 October). 10.1371/journal.pbio.3002332

Croze, M., Wollstein, A., Božičević, V., Živković, D., Stephan, W., & Hutter, S. (2017). A genome-wide scan for genes under balancing selection in Drosophila melanogaster. BMC Evolutionary Biology, 17(1), 1–12. 10.1186/s12862-016-0857-z

Enell, L. E., Kapan, N., Söderberg, J. A. E., Kahsai, L., & Nässe, D. R. (2010). Insulin signaling, lifespan and stress resistance are modulated by metabotropic GABA receptors on insulin producing cells in the brain of Drosophila. PLoS ONE, 5(12). 10.1371/journal.pone.0015780

Fast, I., Hewel, C., Wester, L., Schumacher, J., Gebert, D., Zischler, H., Berger, C., & Rosenkranz, D. (2017). Temperature-responsive miRNAs in Drosophila orchestrate adaptation to different ambient temperatures. Rna, 23(9), 1352–1364. 10.1261/rna.061119.117

Feller, C., Forné, I., Imhof, A., & Becker, P. B. (2015). Global and specific responses of the histone acetylome to systematic perturbation. Molecular Cell, 57(3), 559–571. 10.1016/j.molcel.2014.12.008

Goh, G. H., Blache, D., Mark, P. J., Jason Kennington, W., & Maloney, S. K. (2021). Daily temperature cycles prolong lifespan and have sex-specific effects on peripheral clock gene expression in Drosophila melanogaster. Journal of Experimental Biology, 224(10). 10.1242/jeb.233213

Hostrup, M., Lemminger, A. K., Stocks, B., Gonzalez-Franquesa, A., Larsen, J. K., Quesada, J. P., Thomassen, M., Weinert, B. T., Bangsbo, J., & Deshmukh, A. S. (2022). High-intensity interval training remodels the proteome and acetylome of human skeletal muscle. ELife, 11. 10.7554/eLife.69802

Jang, T., & Lee, K. P. (2018). Context-dependent effects of temperature on starvation resistance in Drosophila melanogaster: Mechanisms and ecological implications. Journal of Insect Physiology, 110, 6–12. 10.1016/j.jinsphys.2018.08.004

Johnston, I. A., & Dunn, J. (1987). Temperature acclimation and metabolism in ectotherms with particular reference to teleost fish. Symposia of the Society for Experimental Biology, 41(May), 67–93.

Karan, D., & David, J. R. (2000). Cold tolerance in Drosophila: Adaptive variations revealed by the analysis of starvation survival reaction norms. Journal of Thermal Biology, 25(5), 345–351. 10.1016/S0306-4565(99)00106-0

Klepsatel, P., Wildridge, D., & Gáliková, M. (2019). Temperature induces changes in Drosophila energy stores. Scientific Reports, 9(1), 1–10. 10.1038/s41598-019-41754-5

Kosar, F., Akram, N. A., Sadiq, M., Al-Qurainy, F., & Ashraf, M. (2019). Trehalose: A Key Organic Osmolyte Effectively Involved in Plant Abiotic Stress Tolerance. Journal of Plant Growth Regulation, 38(2), 606–618. 10.1007/s00344-018-9876-x

Koyama, T., Texada, M. J., Halberg, K. A., & Rewitz, K. (2020). Metabolism and growth adaptation to environmental conditions in Drosophila. Cellular and Molecular Life Sciences, 77(22), 4523–4551. 10.1007/s00018-020-03547-2

Levine, M. T., & Begun, D. J. (2008). Evidence of spatially varying selection acting on four chromatin-remodeling loci in Drosophila melanogaster. Genetics, 179(1), 475–485. 10.1534/genetics.107.085423

Lin, Y. C., Zhang, M. Y., Chang, Y. J., & Kuo, T. H. (2023). Comparisons of lifespan and stress resistance between sexes in Drosophila melanogaster. Heliyon, 9(8), e18178. 10.1016/j.heliyon.2023.e18178

Love, M. I., Huber, W., & Anders, S. (2014). Moderated estimation of fold change and dispersion for RNA-seq data with DESeq2. Genome Biology, 15(12), 1–21. 10.1186/s13059-014-0550-8

Mołoń, M., Dampc, J., Kula-Maximenko, M., Zebrowski, J., Mołoń, A., Dobler, R., Durak, R., & Skoczowski, A. (2020). Effects of temperature on lifespan of drosophila melanogaster from different genetic backgrounds: Links between metabolic rate and longevity. Insects, 11(8), 1–18. 10.3390/insects11080470

Nakagawa, T., Ikehara, T., Doiguchi, M., Imamura, Y., Higashi, M., Yoneda, M., & Ito, T. (2015). Enhancer of acetyltransferase chameau (EAChm) is a novel transcriptional co-activator. PLoS ONE, 10(11), 1–14. 10.1371/journal.pone.0142305

Padmanabha, H., Lord, C. C., & Lounibos, L. P. (2011). Temperature induces trade-offs between development and starvation resistance in Aedes aegypti (L.) larvae. Medical and Veterinary Entomology, 25(4), 445–453. 10.1111/j.1365-2915.2011.00950.x

Peleg, S., Feller, C., Forne, I., Schiller, E., Sévin, D. C., Schauer, T., Regnard, C., Straub, T., Prestel, M., Klima, C., Schmitt Nogueira, M., Becker, L., Klopstock, T., Sauer, U., Becker, P. B., Imhof, A., & Ladurner, A. G. (2016). Life span extension by targeting a link between metabolism and histone acetylation in Drosophila . EMBO Reports, 17(3), 455–469. 10.15252/embr.201541132

Pijpe, J., Brakefield, P. M., & Zwaan, B. J. (2007). Phenotypic plasticity of starvation resistance in the butterfly Bicyclus anynana. Evolutionary Ecology, 21(5), 589–600. 10.1007/s10682-006-9137-5

Pino, L. K., Searle, B. C., Bollinger, J. G., Nunn, B., MacLean, B., & MacCoss, M. J. (2020). The Skyline ecosystem: Informatics for quantitative mass spectrometry proteomics. Mass Spectrometry Reviews, 39(3), 229–244. 10.1002/mas.21540

Rauschenbach, I. Y., Karpova, E. K., Adonyeva, N. V., Andreenkova, O. V., Faddeeva, N. V., Burdina, E. V., Alekseev, A. A., Menshanov, P. N., & Gruntenko, N. E. (2014). Disruption of insulin signalling affects the neuroendocrine stress reaction in drosophila females. Journal of Experimental Biology, 217(20), 3733–3741. 10.1242/jeb.106815

Ritchie, M. E., Phipson, B., Wu, D., Hu, Y., Law, C. W., Shi, W., & Smyth, G. K. (2015). Limma powers differential expression analyses for RNA-sequencing and microarray studies. Nucleic Acids Research, 43(7), e47. 10.1093/nar/gkv007

Ronowska, A., Szutowicz, A., Bielarczyk, H., Gul-hinc, S., Zy, M., & Jankowska-kulawy, A. (2018). The Regulatory Effects of Acetyl-CoA Distribution in the Healthy and Diseased Brain. Frontiers in Cellular Neuroscience, 12(July), 1–20. 10.3389/fncel.2018.00169

Santos, J. L., Nick, F., Adhitama, N., Fields, P. D., & Stillman, J. H. (2024). Trehalose mediates salinity-stress tolerance in a crustacean. BioRxiv. 10.1101/2024.02.09.579427

Shell, B. C., Schmitt, R. E., Lee, K. M., Johnson, J. C., Chung, B. Y., Pletcher, S. D., & Grotewiel, M. (2018). Measurement of solid food intake in Drosophila via consumption-excretion of a dye tracer. Scientific Reports, 8(1), 1–13. 10.1038/s41598-018-29813-9

Sudhakar, S. R., Pathak, H., Rehman, N., Fernandes, J., Vishnu, S., & Varghese, J. (2020). Insulin signalling elicits hunger-induced feeding in Drosophila. Developmental Biology, 459(2), 87–99. 10.1016/j.ydbio.2019.11.013

Tan, D., Wei, W., Han, Z., Ren, X., Yan, C., Qi, S., Song, X., Zheng, Y. G., Wong, J., & Huang, H. (2022). HBO1 catalyzes lysine benzoylation in mammalian cells. IScience, 25(11), 1–13. 10.1016/j.isci.2022.105443

Tellis, M. B., Kotkar, H. M., & Joshi, R. S. (2023). Regulation of trehalose metabolism in insects: from genes to the metabolite window. Glycobiology, February, 262–273. 10.1093/glycob/cwad011

Tennessen, J. M., Barry, W. E., Cox, J., & Thummel, C. S. (2014). Methods for studying metabolism in Drosophila. Methods, 68(1), 105–115. 10.1016/j.ymeth.2014.02.034

Thorat, L. J., Gaikwad, S. M., & Nath, B. B. (2012). Trehalose as an indicator of desiccation stress in Drosophila melanogaster larvae: A potential marker of anhydrobiosis. Biochemical and Biophysical Research Communications, 419(4), 638– 642. 10.1016/j.bbrc.2012.02.065

Umezaki, Y., Hayley, S. E., Chu, M. L., Seo, H. W., Shah, P., & Hamada, F. N. (2018). Feeding-State-Dependent Modulation of Temperature Preference Requires Insulin Signaling in Drosophila Warm-Sensing Neurons. Current Biology, 28(5), 779–787.e3. 10.1016/j.cub.2018.01.060

Venkatasubramani, A. V., Ichinose, T., Kanno, M., Forne, I., Tanimoto, H., Peleg, S., & Imhof, A. (2023). The fruit fly acetyltransferase chameau promotes starvation resilience at the expense of longevity. EMBO Reports, 1–15. 10.15252/embr.202357023

Wu, H., Moshkina, N., Min, J., Zeng, H., Joshua, J., Zhou, M., & Plotnikov, N. A. (2012). Structural basis for substrate speci fi city and catalysis of human histone acetyltransferase 1. Proceedings of the National Academy of Sciences, 109(23), 8925– 8930. 10.1073/pnas.1114117109

Xiao, Y., Hsiao, T. H., Suresh, U., Chen, H. I. H., Wu, X., Wolf, S. E., & Chen, Y. (2014). A novel significance score for gene selection and ranking. Bioinformatics, 30(6), 801–807. 10.1093/bioinformatics/btr671

Xu, J., Sheng, Z., & Palli, S. R. (2013). Juvenile Hormone and Insulin Regulate Trehalose Homeostasis in the Red Flour Beetle, Tribolium castaneum. PLoS Genetics, 9(6). 10.1371/journal.pgen.1003535

Yamauchi, T., Stegeman, J., & Goldberg, E. (1975). The Effects of Starvation Pentose Phosphate and Temperature Trout Liver l in Brook on Pathway Dehydrogenases The Effects of Temperature and Feeding. Archives of Biochemistry and Biophysics, 167, 13–20.

Yu, G., Wang, L. G., Han, Y., & He, Q. Y. (2012). ClusterProfiler: An R package for comparing biological themes among gene clusters. OMICS A Journal of Integrative Biology, 16(5), 284–287. 10.1089/omi.2011.0118

